# Locomotion selectively enhances visual speed encoding in mouse medial higher visual areas

**DOI:** 10.1101/2025.02.21.639472

**Authors:** Edward A. B. Horrocks, Aman B. Saleem

## Abstract

Mammalian visual systems are comprised of multiple brain areas with distinct functional roles. Whilst functional specialisations have been proposed in the mouse based on visual feature encoding, the extent to which these specialisations are contingent on ongoing behaviour is unknown. To address this we analysed neural encoding of visual motion stimuli by thousands of neurons recorded across six cortical and two thalamic mouse visual areas while mice were stationary or locomoting. We found locomotion selectively enhanced visual speed encoding in medial higher visual cortical areas, indicating that these areas may be specialised for the processing visual motion during locomotion. By contrast, encoding of drifting gratings direction was enhanced non-selectively across the mouse visual cortex during locomotion. Our results therefore reveal how a complex interplay of sensory input and ongoing behaviour differentially shapes the efficacy of sensory processing in mouse higher visual areas.

## Introduction

Mammalian visual systems are comprised of multiple brain areas that are specialised for different functional roles (Van Essen, 1979; Wang and Burkhalter, 2007). In primates, two parallel cortical visual processing streams have been identified - a ventral stream associated with object recognition and discrimination and a dorsal stream associated with encoding the spatial location and visual motion of objects, as well as enabling visually-guided actions such as locomotion including through the encoding of optic flow (Goodale and Milner, 1992; Van Essen and Gallant, 1994; Andersen, 1997; Milner and Goodale, 2006; Pisella et al., 2006).

Analogous dorsal and ventral cortical processing streams have been proposed in the mouse visual system. Mouse dorsal and ventral streams were first proposed on the basis of anatomical connectivity. Areas within putative dorsal and ventral streams exhibit stronger within-stream connectivity as well as differences in their downstream connections ventral stream areas preferentially connect to temporal and parahippocampal areas associated with object recognition whereas dorsal stream areas preferentially connect to motor, parietal and limbic cortex (Wang, Gao and Burkhalter, 2011; Wang, Sporns and Burkhalter, 2012; Glickfeld et al., 2013; Wang and Burkhalter, 2013). The functional response properties of ventral and dorsal stream areas have also been investigated. Neurons in ventral stream areas tend to exhibit response properties suited to texture and object discrimination whilst dorsal stream areas have properties generally more suited to the encoding of visual motion (Marshel et al., 2011; Murakami, Matsui and Ohki, 2017; Nishio et al., 2018; Han, Vermaercke and Bonin, 2022; Yu et al., 2022),

Sensory processing in the mouse is strongly modulated by behaviour (Busse et al., 2017; Parker et al., 2020; Horrocks, Mareschal and Saleem, 2022). In particular, locomotion has strong effects on visual processing, including enhancing the encoding of visual speed in mouse primary visual cortex (Horrocks, Rodrigues and Saleem, 2024). A key outstanding question is whether such effects of behaviour on sensory processing vary between visual areas. Selective enhancement of specific visual feature encoding during different behaviours would provide important clues as to the functional specialisations of mouse visual areas. Notably, neurons in different mouse visual areas exhibit varying tuning properties for locomotion speed (Saleem et al., 2013; Erisken et al., 2014; Blot et al., 2021; Christensen and Pillow, 2022), yet despite these differences the limited available evidence suggests that locomotion influences visual feature encoding similarly across mouse cortical visual areas (Christensen and Pillow, 2022). Here, we investigated whether visual speed encoding, a feature associated with dorsal-stream processing, is selectively enhanced in specific mouse visual areas during locomotion.

We leveraged large-scale in vivo extracellular electrophysiological recordings of thousands of neurons in six cortical and two thalamic mouse visual areas (Allen ‘Visual Coding’ dataset; (Siegle et al., 2021)) to analyse the encoding of visual speed during stationary and locomotion behavioural states. We first characterised visual speed tuning properties and found that they differed substantially between visual areas. Specifically, broad ranges of tuning properties in V1 and thalamic areas were consistent with functional roles distributing varied information about visual motion throughout the mouse visual system. Narrower ranges of tuning properties in higher cortical visual areas were suggestive of more specialised roles in the processing of visual motion, with a posterior-to-anterior gradient of faster preferred visual speeds. When we compared visual speed encoding between stationary and locomotion states we found that locomotion selectively enhanced visual speed encoding in medial higher visual areas AM and PM. In contrast, locomotion non-selectively enhanced drifting gratings direction encoding across mouse visual cortex, indicating that the selective enhancement of medial higher visual areas during locomotion was specific to the encoding of visual speed. These findings suggest that medial higher visual areas, which are biased to process the peripheral visual field, may be specialised for the encoding of locomotion-related optic flow. More generally, our findings establish that the effects of behaviour on sensory processing exhibit a complex brain area-dependence that is contingent upon the specific sensory feature being encoded.

## Results

We investigated the encoding of visual speed across across the mouse visual system during different behavioural states by analysing the firing rate responses of thousands of neurons (n=5,707) to moving dot field stimuli whilst mice were in stationary or locomoting states (n = 19 mice; (Siegle et al., 2021)). In this dataset, individual mice tended to either locomote or remain stationary within a recording session. Six cortical visual areas (V1, LM, AL, RL, AM, PM) and two thalamic visual areas (LGN, LP) were simultaneously targeted in each recording using 6 neuropixel probes. Stimuli consisted of fields of white dots which covered a large proportion of the contralateral visual field (120º azimuth and 95º elevation) and moved at one of seven visual speeds (0, 16, 32, 64, 128, 256, 512 º/s). We refer to single neuron firing rate responses as a function of these visual speeds as visual speed tuning.

### Four classes of visual speed tuning across the mouse visual system

We discovered four classes of visual speed tuning curve in the mouse visual system (Figure 1). To characterise the shapes of tuning curves from neurons across the six cortical and two thalamic areas recorded we performed a hierarchical clustering procedure on all reliable tuning curves (n= 4,706; Supp. Figure 1a), where reliability was assessed using cross-validated coefficient of determination (Horrocks, Rodrigues and Saleem, 2024). This revealed that visual speed tuning curves could broadly be classified into four distinct shapes: lowpass, bandpass, bandreject and highpass (Supp. Figure 1b). To characterise the encoding properties of these different tuning classes we then classified each reliable tuning curve based on the best-fitting of four representative descriptive functions (Figure 1c,d). The majority of tuned cells exhibited bandpass filtering properties for visual speeds presented (63%; Figure 1e). Lowpass and bandreject were the next most frequent tuning shapes (lowpass: 15%, bandreject: 14%) followed by highpass (8%).

**Figure 1:**
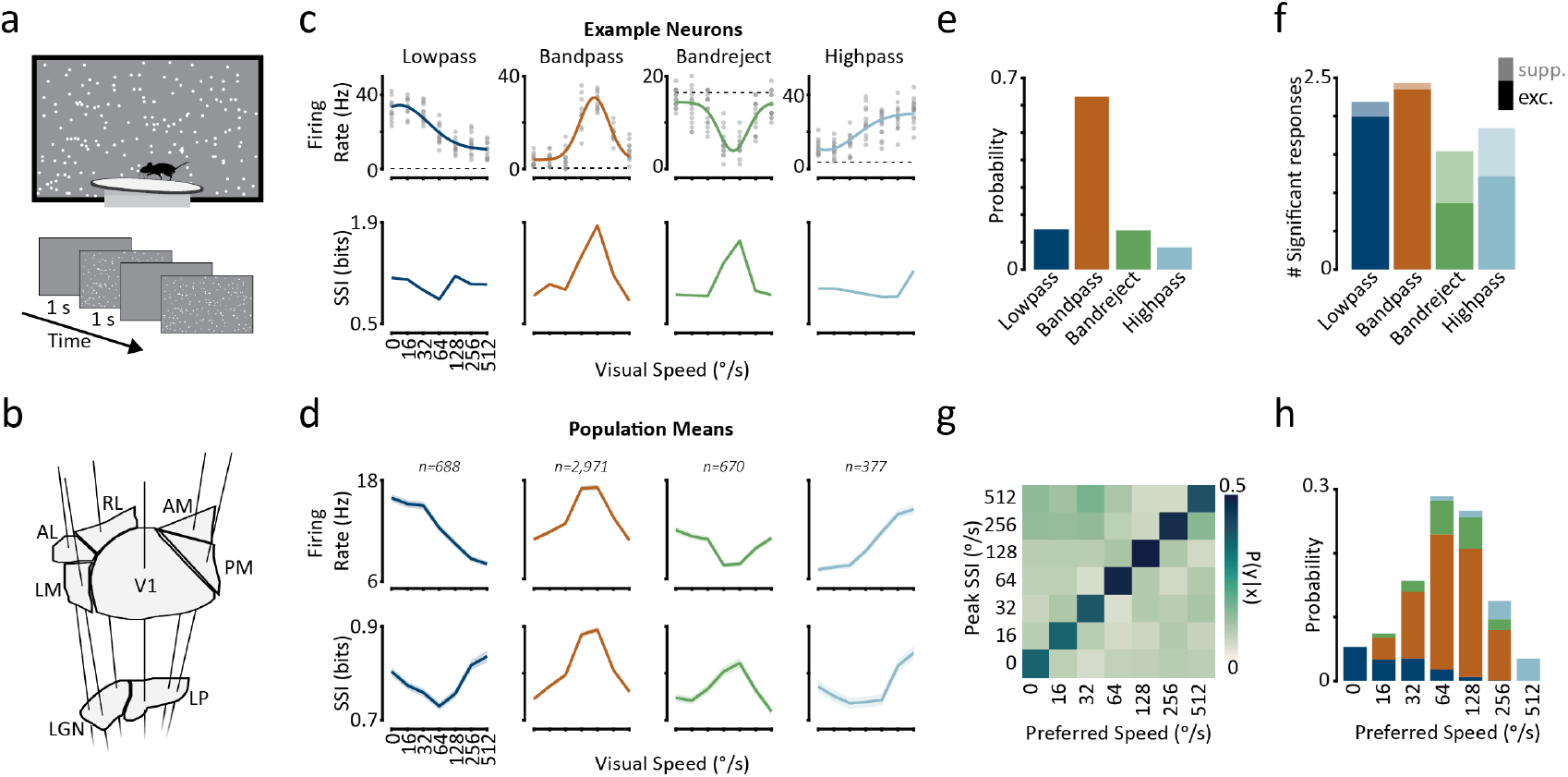
Four classes of visual speed tuning curve across the mouse visual system. **a** Schematic of experimental setup. Mice were presented trials of moving random dot field stimuli whilst free to locomote on a circular treadmill. **b** Schematic of neuropixel probe insertions - six probes simultaneously targeted six cortical and two thalamic visual areas. **c** Examples of four different tuning curve classes. Top row: Example tuning curves. Each grey circle is a single-trial spike count. Thicker lines are gaussian descriptive function fits. Dashed lines indicate mean pre-stimulus baseline firing rates. Bottom row: Stimulus-specific information (SSI) for the corresponding examples in the top row. **d** Same as **a** for population means of each tuning class. Shaded regions indicate the mean ± SEM. Numbers above plots indicate the number of classified tuning curves within each class. **e** Probability histogram of tuning class for all reliable tuning curves. **f** Stacked bar chart showing the number of significant responses (sign-rank test of evoked firing rate vs baseline firing rate) for each tuning class. Bars are split into excitatory (dark colours) and suppressed (light colours) responses, based on the difference from activity during pre-stimulus baseline periods. **g** Discrete conditional probability distribution of P(Peak SSI | Preferred Speed) showing correspondence between the two measures. **h** Probability histograms of preferred visual speed for all tuning curves. Stacked bars are colour code according to the classified tuning shape in **c-f**.

Excitation and suppression differentially shape visual speed tuning curve classes. To understand whether different tuning curves we observed were related to excitation or suppression, we compared the stimulus responses to pre-stimulus baseline activity levels. Bandpass and lowpass tuning curves were shaped primarily by selective excitation to specific visual speeds (Figure 1f). By contrast, highpass and bandreject tuning shapes were shaped by a mix of selective excitation and suppression (Figure 1f).

Different tuning curve classes preferentially encode different visual speeds. To characterise how well different visual speeds are encoded by different tuning curve classes we used stimulus specific information (SSI), a mutual information (Shannon, 1948) measure that quantifies how much information is present in responses to a specific stimulus (Butts and Goldman, 2006). For example, a neuron that only responds above baseline levels to one stimulus will have high SSI for that stimulus and low SSI for all others. SSI tended to be positively correlated with spike counts for bandpass and highpass tuning classes and negatively correlated for the bandreject class (Figure 1c, d). For the lowpass tuning class SSI peaked for both the slowest and fastest speeds i.e. the stimuli that evoked the largest and smallest spike count responses. Overall, visual speeds that evoked the maximum SSI response generally corresponded to the preferred visual speed of a neuron (Fig 1g, h). Thus, neurons with low-pass tuning curves best encode slow and fast visual speeds, neurons with highpass tuning curves best encode fast visual speeds and neurons with bandpass or bandreject tuning curves best encode intermediate visual speeds.

Distributions of visual speed tuning class vary between mouse visual areas. Having characterised visual speed tuning across the mouse visual system, we next investigated how it varies between areas. Whilst bandpass tuning was the most common class of tuning curve in each visual area (range: 45 - 74%), the overall distribution of tuning classes varied between areas (Figure 2b). For example, in V1 and LP, 25% of tuning curves were classified as lowpass whereas this dropped to 7% for RL and AM, with other areas exhibiting intermediate proportions of lowpass neurons. Interestingly, bandreject tuning comprised the second most common tuning class in areas AL (14%), RL (19%), AM (15%), PM (15%) and LGN (22%), demonstrating that this tuning curve shape is widespread in the mouse visual system and in turn that selective suppression plays an important role in shaping visual speed tuning (Figure 1f).

**Figure 2:**
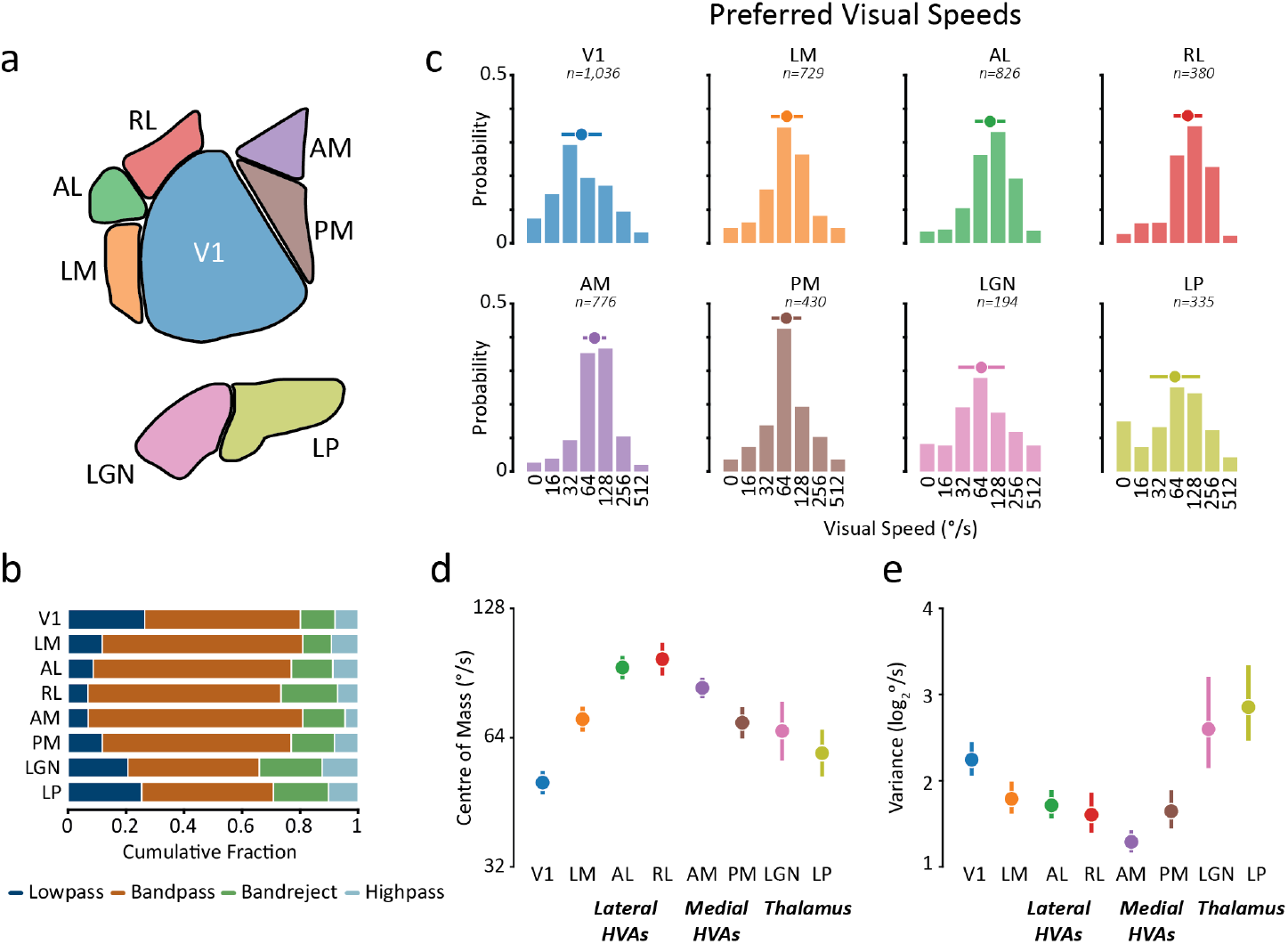
Visual speed tuning properties vary between mouse visual areas. **a** Schematic of 8 mouse visual areas (top - cortex, bottom - thalamus) illustrating the colour code applied to all figure panels. **b** Distributions of tuning class for each visual area. **c** Distributions of preferred visual speeds for each visual area. Circles and horizontal lines indicate the centre of mass and variance of each distribution. Numbers above plots indicate the number of contributing tuning curves. **d** Centre of mass of preferred visual speeds distributions for each brain area (errorbars indicate means and 95% confidence intervals). **e** Same as **d** for the variance of preferred visual speeds distributions.

Distributions of preferred visual speeds vary between mouse visual areas. All higher visual cortical areas had faster speed preferences than V1 on average (Fig 2c,d). Amongst higher visual areas there was an anterior-to-posterior gradient of fast-to-slow speed preferences, with anterior areas AL, RL and AM having the fastest speed preferences and LM and PM the slowest. Thalamic nuclei had speed preferences intermediate between V1 and higher visual cortical areas. The variance of preferred visual speed distributions also varied between visual areas (Fig 2e). V1 and thalamic nuclei LGN and LP had broad distributions of visual speed preferences, consistent with the idea that projection neurons from these areas convey diverse information about visual speed to other visual areas (Glickfeld et al., 2013; Oh et al., 2014; Tohmi et al., 2014; Roth et al., 2016; Blot et al., 2021). Higher visual cortical areas in contrast had more concentrated distributions of visual speed preferences, suggesting that these areas may be specialised for encoding specific ranges of visual speeds.

### Locomotion selectively enhances visual speed tuning in medial higher visual areas AM and PM, as well as V1 and LGN

We next investigated whether the prevalence and strength of visual speed tuning varied between behavioural states in different visual areas. We assessed the strength of tuning using the cross-validated coefficient of determination method (R2; Figure 3a; (Horrocks, Rodrigues and Saleem, 2024), a metric that determines how reliable a tuning curve is across repeated trials compared to a mean firing rate model. We classified a neuron to be tuned for visual speed if its tuning strength was significant (assessed using a shuffled distribution of trial spike counts) and greater than a threshold (*R*2 *≥* 0.1). We obtained similar results regardless of the specific tuning strength threshold we used (Supp. Figure 2).

**Figure 3:**
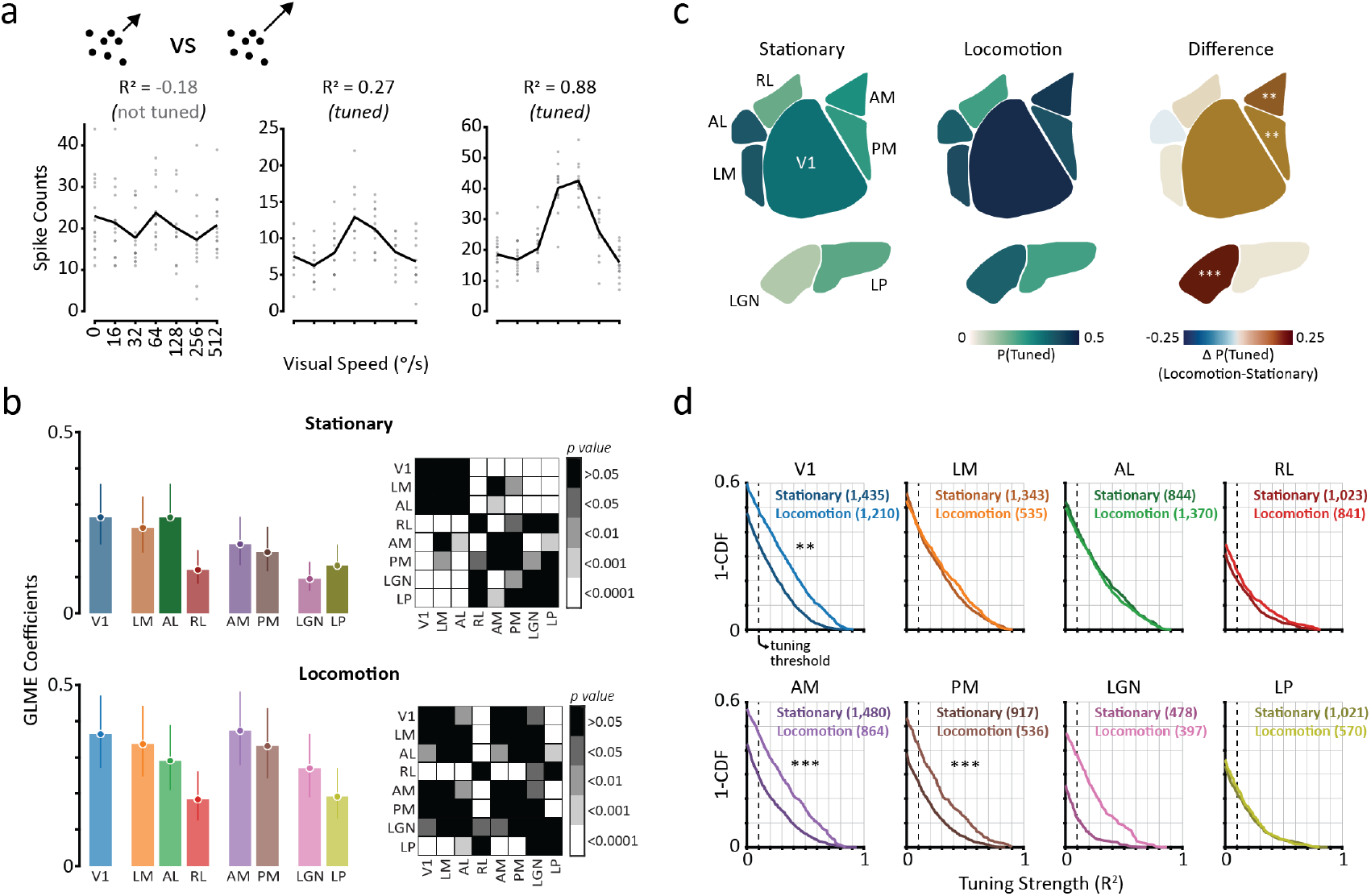
Locomotion selectively enhances visual speed tuning in medial higher visual areas, as well as V1 and LGN. **a** Example tuning curves with different tuning strength (R2) values. Circles indicate individual trial spike counts and black lines the stimulus-conditioned mean of those spike counts. Inset icon illustrates stimulus consisting of fields dots moving dots. **b** GLME model analysis for the probability of visual speed tuning in each visual area during stationary and locomotion states. Bar charts show model coefficients for each area during stationary (top) and locomotion (bottom) states. Errorbars indicate mean ± SEM. Matrices (right) show pairwise comparisons of visual areas for the corresponding states. p-values are Bonferroni corrected. **c** Probability of tuning for visual speed by visual area during stationary states (left), locomotion states (centre) and the difference between locomotion and stationary states (right). ** *p<0*.*01*, \*\*\**p<0*.*001* (Bonferroni-corrected) GLME statistical tests for effect of state on probability of being tuned, for each visual area. **d** Distributions of tuning strength for each visual area, separately for stationary (dark colours) and locomotion (light colours) states. Numbers in brackets indicate number of contributing tuning curves. ** *p<0*.*01*, \*\**\*p<0*.*001* (Bonferroni-corrected).

The prevalence of visual speed tuning varied between visual areas during stationary states (Figure 3 b,c). Lateral cortical visual areas AL and LM had the highest prevalence of visual speed tuned cells (both 38%; Figure 3b), followed by V1 (33%), AM (27%) and PM (24%). Area RL had the lowest prevalence of visual speed tuning (19%) of the cortical areas. The prevalence of tuning was lower in thalamic areas (LGN: 11%, LP: 20%) compared to cortex (Figure 3b).

The weak tuning in LGN suggests that V1 either integrates weakly tuned feedforward inputs from LGN or recurrent cortical circuits strengthen visual speed tuning in V1 during this state.

Locomotion selectively enhanced tuning for visual speed in medial higher visual areas AM and PM, as well as V1 and LGN. When comparing the prevalence and strength of visual speed tuning between stationary and locomotion states we found clear area-dependent effects. Specifically, there was a significant increase in the prevalence of visual speed tuning in medial higher visual areas AM (*p<0*.*01*) and PM (*p<0*.*01*) during locomotion, but no significant change in lateral higher visual areas (Figure 3c). Similarly, distributions of tuning strength shifted towards higher values for AM (*p<0*.*001*, LME model analysis) and PM (*p<0*.*001*), but not for lateral higher visual areas during locomotion (Figure 3d). We also observed an increase in the strength of tuning for neurons in V1 during locomotion (*p<0*.*01*), in agreement with our previous findings (Horrocks, Rodrigues and Saleem, 2024), as well as an increase in the prevalence of tuning in LGN (*p<0*.*001*). As a result of these changes, AM and V1 had the strongest tuning for visual speed during locomotion, compared to AL and LM in stationary sessions (Supp. Figure 2).

### Locomotion selectively enhances visual speed population decoding in medial higher visual areas AM and PM, as well as V1 and LGN

We next investigated how locomotion affects visual speed decoding at a population level. Stimulus decoding can provide a measure of how much stimulus information is contained with the responses of populations of neurons. We therefore compared decoding performance of populations of neurons between behavioural states, for each visual area (Figure 4), using an independent Bayes decoder (Zhang et al., 1998; Jazayeri and Movshon, 2006; Graf et al., 2011). We tested small populations of simultaneously recorded neurons (n=10) at a time to facilitate a comparison of the different visual areas and behavioural states.

**Figure 4:**
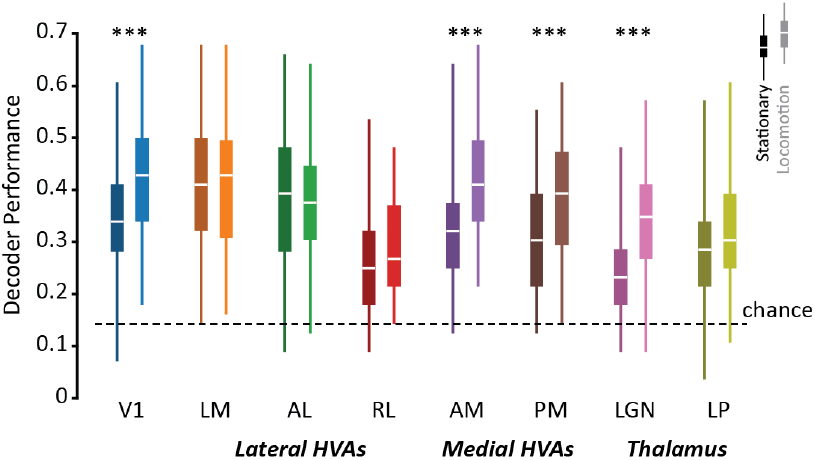
Locomotion selectively enhances visual speed population decoding in medial higher visual areas AM and PM, as well as V1 and LGN. Naive bayes decoding performance (fraction correctly predicted trials) of subsampled populations (n=10) in each visual area during stationary (darker left-side boxplots) and locomoting (lighter right-side box plots) states. Box plots indicate full distributions (boxes are interquartile range, white lines are medians). *** *p<0*.*001* (Bonferroni-corrected) GLME statistical tests for effect of state on decoding performance, for each visual area.

Locomotion selectively enhanced population decoding of visual speed in medial higher visual areas AM and PM, as well as V1 and LGN. We found that locomotion was associated with enhanced decoding of visual speed in medial higher visual areas AM (mean ± SEM performance during stationary states: 0.32 ± 0.01, locomotion states: 0.41 ± 0.01, *p<0*.*001* effect of state, LME model analysis) and PM (stationary: 0.30 ± 0.01, locomotion: 0.38 ± 0.02, *p<0*.*001*) as well as V1 (stationary: 0.34 ± 0.01, locomotion: 0.43 ± 0.01, *p<0*.*001*) and LGN (stationary: 0.24 ± 0.01, locomotion: 0.32 ± 0.01, *p<0*.*001*), in accordance with our findings of improved visual speed tuning in these areas. By contrast, we found no significant difference in decoding performance between states for the lateral higher visual areas (LM, AL and RL) or the thalamic nucleus LP (all *p>0*.*05*). Thus, the area-specific changes in visual speed tuning we observed during locomotion resulted in area-specific changes in the ability to decode visual speed from neural population activity.

### Locomotion non-selectively enhances drifting gratings direction tuning across mouse visual cortex

Does the selective enhancement of medial higher visual areas AM and PM during locomotion reflect general changes in visual processing across visual areas, or are these changes stimulus-specific? To address this question, we repeated our visual feature tuning analysis for drifting gratings direction (Figure 5) in stationary and locomotion states using a related dataset from Allen Institute (‘Brain Observatory 1.1’ stimulus set; Siegle et al., 2021).

**Figure 5:**
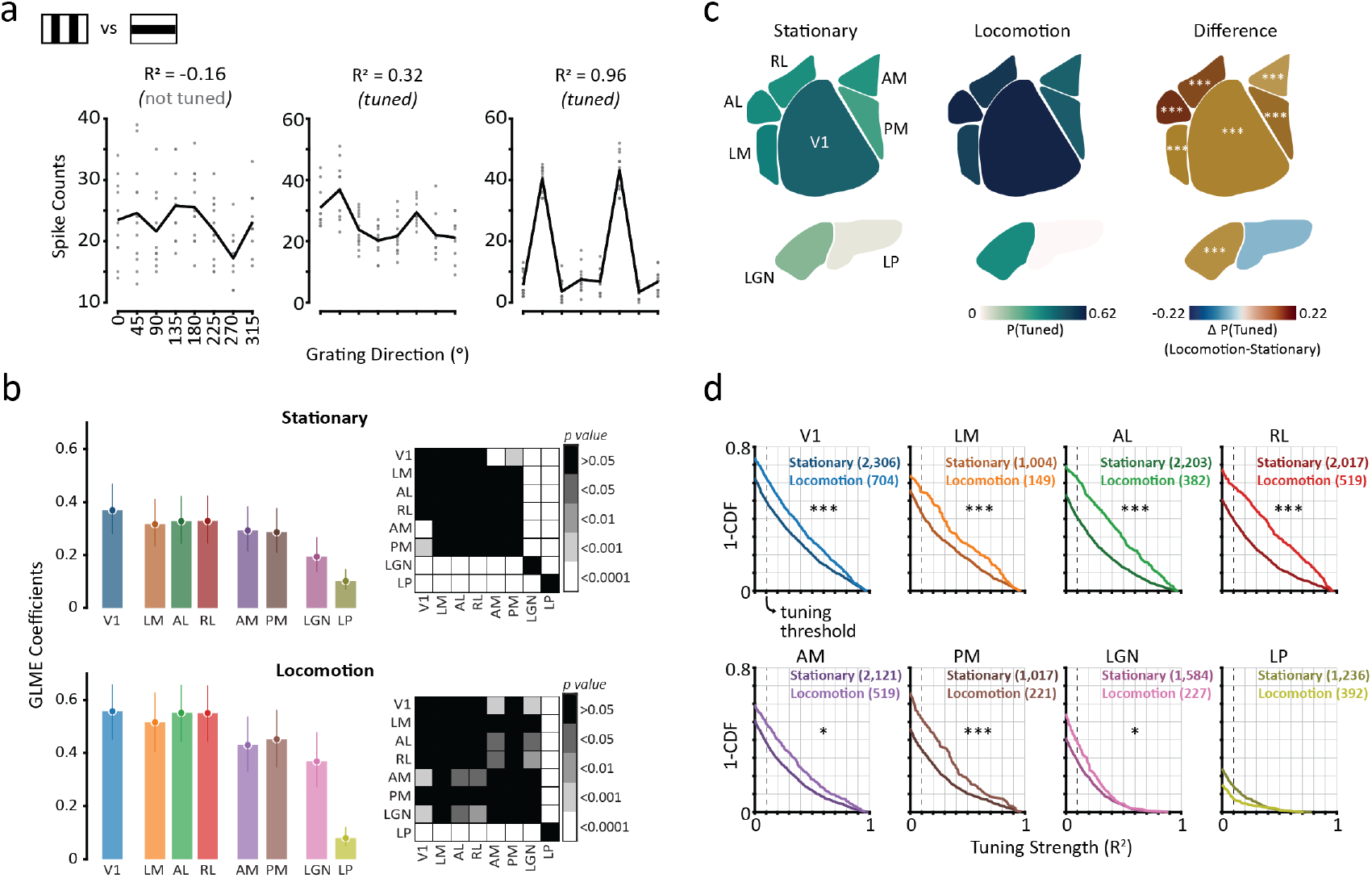
Locomotion non-selectively enhances drifting gratings direction tuning across mouse visual cortex. **a** Example tuning curves with different tuning strength (R2) values. Circles indicate individual trial spike counts and black lines the stimulus condition mean of those spike counts. Inset icon illustrates stimulus consisting of drifting gratings. **b** GLME model analysis for the probability of visual speed tuning in each visual area during stationary and locomotion states. Bar charts show model coefficients for each area during stationary (top) and locomotion (bottom) states. Errobars indicate mean ± SEM. Matrices (right) show pairwise comparisons of visual areas for the corresponding states. p-values are bonferroni corrected. **c** Probability of tuning for drifting grating direction by visual area during stationary states (left), locomotion states(centre) and the difference between locomotion and stationary states (right). *** *p<0*.*001* (Bonferroni-corrected) GLME statistical tests for effect of state on probability of being tuned, for each visual area. **d** Distributions of tuning strength for each visual area, separately for stationary (dark colours) and locomotion (light colours) states. Numbers in brackets indicate number of contributing tuning curves. ** *p<0*.*01*, \*\*\**p<0*.*001* (Bonferroni-corrected).

Locomotion non-selectively enhanced drifting gratings direction tuning across all cortical areas. In contrast to the encoding of visual speed, there was no selective change in the encoding of drifting gratings direction during locomotion (Figure 5b-d). Instead both the prevalence and strength of direction tuning increased for all visual areas during locomotion, with the exception of LP. Indeed, the largest increases in the prevalence of drifting gratings direction tuning were in lateral visual areas AL and RL, in contrast to our findings of selective increases in the prevalence of visual speed tuning in medial higher visual areas AM and PM. Notably, neurons in area RL were robustly tuned for drifting gratings direction in both behavioural states (Figure 5), in contrast to our findings of weak visual speed tuning (Figure 3), indicating that moving dot fields are particularly ill-suited to driving neurons in area RL (see also Sit and Goard, 2020). Our results therefore establish that the effects of behaviour on sensory processing exhibit a complex brain area-dependence that is contingent upon the specific sensory feature being encoded.

## Discussion

Here, by analysing the neural activity of thousands of neurons we have characterised visual speed tuning across six cortical and two thalamic mouse visual areas during stationary and locomotion behavioural states. Our main finding is that behavioural state can selectively alter the efficacy of visual feature encoding in mouse cortical visual areas, and that this area-dependence is contingent on the stimulus feature being encoded. Specifically, we found that the encoding of visual speed is selectively enhanced in medial higher visual areas AM and PM during locomotion. In contrast, the encoding of drifting grating direction is non-selectively enhanced across all higher visual cortical areas. Our findings therefore reveal how ongoing behaviour shapes the functional specialisations of mouse visual areas.

Why are improvements in visual speed encoding during locomotion specific to medial higher visual areas AM and PM? Both areas have been identified anatomically as mouse dorsal stream areas with connectivity to motor and navigation-related areas (Wang, Sporns and Burkhalter, 2012). Indeed, AM and PM are located between primary visual cortex and retrosplenial cortex, an area considered important for visuospatial processing during navigation and with strong connections to the hippocampus. AM and PM exhibit a number of features that suggest their suitability for the encoding of movement-related optic flow such as large receptive field sizes and weak surround suppression (Murgas et al., 2020; Siegle et al., 2021). Importantly, areas AM and PM also have representations of the visual field that are biased towards the periphery (Garrett et al., 2014; Zhuang et al., 2017), where optic flow vectors are generally larger and therefore likely to be most informative about self-motion (Saleem, 2020). These features are concordant with properties of primate dorsal stream area MST, which is specialised for the encoding of optic flow; neurons in area MST have large receptive field sizes, preferentially receive projections from populations that represent the peripheral visual field and can be modulated by non-visual self-motion signals (Gattas et al., 1997; Gu et al., 2006; Ungerleider et al., 2008). The enhanced encoding of the visual speed in AM and PM during locomotion may therefore reflect a functional specialisation for the encoding of visual flow during self-movement.

The role of mouse higher visual area PM has been particularly ambiguous. Whilst anatomically located within the mouse dorsal stream (Wang, Sporns and Burkhalter, 2012), the spatiotemporal tuning properties of neurons within PM are more ventral stream-like i.e. selectivity for high spatial frequency and low temporal frequency (Andermann et al., 2011; Marshel et al., 2011; Han, Vermaercke and Bonin, 2022). Indeed, in agreement with a more ventral stream-like role, visual speed encoding was poor in PM in stationary states. However, during locomotion there was a clear enhancement of visual speed encoding, suggesting that the functional role of PM, along with other mouse visual areas, is contingent on ongoing behaviour and context more generally (Jin and Glickfeld, 2020).

The bias for superior visual speed encoding in lateral higher visual areas in stationary mice (Figure 3) is unexpected. Whilst area AL has been characterised as a dorsal stream area (Wang, Sporns and Burkhalter, 2012) and has been previously shown to perform well encoding the motion direction of random dot kinematograms similar to those used here (Yu et al., 2022), area LM is often considered a ventral stream area with properties more suitable for visual texture discrimination (Yu et al., 2022; Bolaños et al., 2024). This lateral bias is not a general principle of visual processing, since there was no significant difference in the prevalence of drifting gratings tuning between higher visual cortical areas during stationary states (Figure 5). Nor was this lateral bias for visual speed encoding present during locomotion, instead replaced by a more even distribution of visual speed encoding across the mouse visual cortex (Figure 3b).

Our analysis of how visual speed tuning properties vary between visual areas indicates that the visual field biases of different mouse higher visual areas are a key feature of their functional specialisations. Most strikingly, the variance of preferred visual speed distributions differed substantially between areas, with broad distributions of preferred visual speeds in V1 and thalamic areas LGN and LP. These broad distributions are consistent with these areas distributed varied visual motion information throughout the mouse visual cortex (Blot et al., 2021; Han and Bonin, 2024). The narrower distributions in higher visual cortical areas suggest more specialised roles for processing specific visual speeds. Interestingly, we observed an anatomical gradient of preferred visual speeds, with neurons in anterior higher visual areas AL, RL and AM exhibiting the fastest preferred speeds overall. Anterior higher cortical visual areas are biased to represent lower elevations of the visual field (Zhuang et al., 2017), and so faster visual speed preferences in these areas might reflect faster visual speeds experienced in this part of the visual field owing to the proximity of the floor plane.

Our results reveal important insights into the functional roles of mouse higher visual cortical areas by demonstrating that the effects of behaviour are both brain area and stimulus-dependent. These findings emphasise a complexity of function that is highly context-dependent, establishing the importance of considering behaviour, and context more generally, when determining functional specialisations of mouse visual areas. As such, future experimentation taking into account ongoing behaviour and task context will be important for elaborating the functional roles mouse visual areas serve in visual perception and visually-guided action.

## Ackowledgements

This work was supported by The Sir Henry Dale Fellowship from the Wellcome Trust and Royal Society (200501); the Human Frontier in Science Program (RGY0076/2018), Biotechnology and Biological Sciences Research Council grant (R004765 and BB/W01579X/1), UKRI Frontier Research grants (EP/Y024656/1) to A.B.S.; and Biotechnology and Biological Sciences Research Council studentship to E.H. We thank Sam Solomon for comments on the manuscript.

## Author Contributions

This work was conceptualised by E.H. and A.B.S.; Methodology, software and formal analysis were by E.H.; Visualisation and writing by E.H. and A.B.S.; and Supervision and funding acquisition by A.B.S.

## Methods

### Data collection

We analysed a large-scale in-vivo extracellular electrophysiology dataset (Visual Coding - Neuropixels (Siegle et al., 2021)) made open access by the Allen Institute. In each recording session session (one per mouse) six neuropixel probes were acutely inserted to to record extracellularly in six cortical visual areas: primary visual cortex (V1), lateromedial area (LM), anterolateral area (AL), rostrolateral area (RL), anteromedial area (AM) and posteromedial area (PM), and two thalamic areas: dorsal lateral geniculate nucleus (dLGN) and lateral posterior nucleus (LP). During each session, mice were head-fixed and free to locomote on a circular treadmill whilst passively viewing stimuli, of which we analysed a subset.

#### Experimental subjects

We analysed two datasets with separate stimulus sets. Analysis of visual speed encoding used the *Functional Connectivity* stimulus set and analysis of drifting gratings direction encoding used the *Brain Observatory 1*.*1* stimulus set. We analysed 12 *Functional Connectivity* sessions with sufficient stationary trials (n = 7 male, n = 5 female; age 114-142 days; n = 8 wild-type C57BL/6J, n = 3 Sst-IRES-Cre × Ai32, n = 1 Vip-IRES-Cre × Ai32) and 8 *Functional Connectivity* sessions with sufficient locomotion trials (n = 5 male, n = 3 female; age 108-135 days; n = 4 wild-type C57BL/6J, n = 2 Sst-IRES-Cre × Ai32, n = 2 Vip-IRES-Cre × Ai32). We analysed 16 *Brain Observatory 1*.*1* sessions with sufficient stationary trials (n = 13 male, n = 3 female; age 98-140 days; n = 9 wild-type C57BL/6J, n = 4 Pvalb-IRES-Cre × Ai32, n = 3 Vip-IRES-Cre × Ai32) and 6 *Brain Observatory 1*.*1* sessions with sufficient locomotion trials (n = 4 male, n = 2 female; age 93-122 days; n = 1 wild-type C57BL/6J, n = 4 x Sst-IRES-Cre × Ai32, n = 1 Vip-IRES-Cre × Ai32).

#### Visual stimuli

To investigate the encoding of visual speed we analysed neural responses to moving dot field stimuli (*Functional Connectivity* visual stimulus set). Stimuli consisted of fields of 200 white moving dots (diameter = 3º) which covered a large proportion of the contralateral visual field (120º azimuth and 95º elevation). On each trial, dots moved at one of seven visual speeds (speeds = 0, 16, 32, 64, 128, 256, 512º/s) in one of 4 directions (0º, 45º, 90º, 135º, where 0º = left-to-right), with 90% coherence. Stimulus duration was 1s with a 1s grey screen inter-stimulus-interval.

We also analysed the encoding of drifting grating direction (*Brain Observatory 1*.*1* visual stimulus set). Full-screen sinusoidal drifting gratings (spatial frequency = 0.04 cycles/º, contrast = 80%) moved in one of 8 directions (0-315º, equally spaced) at one of 5 temporal frequencies (1, 2, 4, 8, 15Hz). Stimulus duration was 2s with a 1s grey screen inter-stimulus-interval.

### Data Analysis

#### Spike sorting

Data were spike-sorted using the Allen Institute’s in-house spike sorting pipeline (Siegle et al., 2021) which uses Kilosort2 (Pachitariu et al., 2016) to perform initial spike sorting followed by a number of custom post-processing modules that remove double-counted spikes and noise units and compute a number of waveform and cluster quality metrics. We restricted our analysis to ‘good’ clusters which we defined based on 3 criteria (Horrocks, Rodrigues and Saleem, 2024; Laboratory et al., 2024): 1) Refractory period violations *≤* 10% 2) Amplitude distribution cut-off *≤* 10% and 3) Mean amplitude *≥* 50*μ*V.

#### Classification of trials according to behavioural state

We classified the behavioural state of trials (defined as the stimulus duration epoch) according to the locomotion speed of mice recorded by a rotary encoder. We classified trials as stationary if trial-mean locomotion speed was <0.5cm/s and remained under 3 cm/s for 75% of the trial. We classified trials as locomotion if trial-mean wheel speed was >3cm/s and remained over 0.5cm/s for 75% of the trial. We have previously shown this to be a robust criteria for defining the locomotor state of trials (Horrocks, Rodrigues and Saleem, 2024). To calculate locomotion speed we resampled the rotary encoder data at 100Hz and smoothed it using a gaussian kernel with a standard deviation of 35ms. To investigate visual speed tuning we considered each direction of motion independently. To investigate drifting grating direction tuning we considered each temporal frequency independently. We only analysed responses where there were at least 10 trials for each visual speed/motion direction that were classified with the same behavioural state (stationary or locomotion).

#### Tuning strength

To assess tuning strength we calculated the cross-validated Coefficient of Determination (R^2^). on trialbased spike counts (Horrocks, Rodrigues and Saleem, 2024). To enable a fair comparison we downsampled trial counts to 10 (minimum required for data inclusion). We implemented 3-fold cross-validation by randomly dividing trials into a training set comprising 2/3 of the trials and a test set comprising the remaining 1/3 (equally sampled from each stimulus condition). During each iteration, two models were created using the training data: a tuning curve model (trained model) representing the mean spike count responses to individual stimulus conditions and a null model representing the average spike count across all stimulus conditions. Using the test data, we constructed a test model by calculating the mean spike count for each stimulus condition. To evaluate the performance of the trained model and null model, we calculated the sum-of-squared residuals between each model and the test model. The coefficient of determination (*R*^2^) was then computed using the following equation:

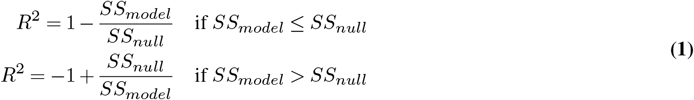

where *SS*_*model*_ is the sum of squared residuals between the *trained model* and the *test model* and *SS*_*null*_ is the sum of squared residuals between the *null model* and the *test model*. We computed the mean *R*^2^ value over the 3 cross-validations, using a unique set of test trials on each iteration. We repeated this entire process 10 times with different random splits of train and test trials, providing 10 estimates of*R*^2^ and took the final estimate of *R*^2^ as the mean of these 10 values. To assess the statistical significance of these tuning strength values we also generated a shuffled distribution of *R*^2^ values for each neuron by performing the same 3-fold cross-validation procedure on randomly shuffled spike counts, repeated 100 times. We considered a neuron to be tuned if *R*^2^ *≥* 0.1. and *R*^2^ *≥* 95th percentile of the shuffled distribution (*p ≥* 0.05). For comparisons of tuning strength we set a floor of *R*^2^ = 0.

#### Tuning curve clustering

To cluster tuning curves for visual speed we used a hierarchical sorting method. We first determined which tuning curves were ‘tuned’ using the criteria above. We then generated a dissimilarity matrix by calculating the euclidean distance between pairs of (z-scored) tuning curves. Following this, we obtained an initial dendrogram (Matlab function linkage) with the unweighted average distance. We then found the optimal leaf order using an algorithm that minimises the sum of pairwise distances between neighbouring leaves (Bar-Joseph, Gifford and Jaakkola, 2001) (Matlab function optimalleaforder). Finally clusters were generated by setting a dendrogram cutoff that produced 8 clusters, which was chosen based on a peak in the mean silhouette score plot for a range of cluster numbers (2:2:64). We obtained similar results for a range of different dendrogram cutoff settings.

#### Tuning shape classification

We classified tuning curves using the best-fitting of four descriptive functions (lowpass, bandpass, bandreject and highpass). Descriptive functions were gaussians paramterised as:

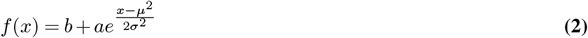

where *b* is a baseline firing rate parameter, *a* is an amplitude parameter, *μ* is the mean of the distribution and *σ* the standard deviation. *x* represents the domain of *log*_2_-transformed stimulus speeds the functions were fitted on and *e* is the Euler’s number. The *log*_2_-transformed value of 0*°/s* was forced to = 3, such that the *log*_2_-transformed stimulus speeds were defined by the vector 3:1:9.

The different tuning classes were differentiated using different parameter bounds. Lowpass and highpass were described by two functions each (with negative and positive amplitudes). Each function has an appropriately bounded mean parameter. The upper bound of bandpass and bandreject was limited to ensure well-defined maxima or minima. Additionally, bandpass and bandreject classifications required fitted functions to have a peak with a prominence of 1/3 of the spike-count range of the tuning curve (assessed using the Matlab function findpeaks). Preferred visual speed classification Preferred visual speeds were classified as the visual speed that evoked the maximum mean spike count response for lowpass, bandpass and highpass tuning curves and as the visual speed that evoked the minimum mean spike count response for bandreject tuning curves.

#### Stimulus-specific (SSI)

To calculate SSI we first first binned responses into 7 quantiles (after downsampling to 10 trials for each stimulus condition) to enable a fair comparison between cells with large differences in firing rates. We then calculated SSI using the following equation from Butts, (2003):

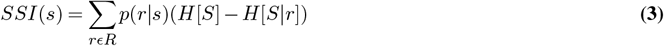

where *SSI*(*s*) is the stimulus specific information for stimulus *s, r* is the set of responses to a given stimulus speed, *S* is the set of all stimulus speeds and *R* is the set of all responses, 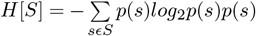 is the entropy of the set of visual speeds, and 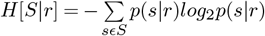 is the entropy of a visual speed given a particular response.

#### Response excitation and suppression

To determine whether a spike-count response to an individual stimulus condition was excitatory or suppressed we compared it to the baseline firing rate (200ms before stimulus onset). For each tuning curve we tested whether there was a significant change in firing rate between pre-stimulus and stimulus epochs using the Bonferroni-corrected Wilcoxon signed-rank test. We then determined responses with significant changes in firing rate as excitatory or suppressed based on the direction of change of firing rate.

#### Statistical analysis of tuning strength

To test whether there were statistically significant differences in the probability of neurons being tuned for visual speed or drifting grating direction between different brain areas and behavioural states we used a generalized linear mixed effects (GLME) model:

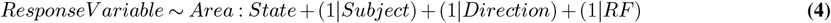

Where *ResponseV ariable* was *P* (*Tuned*), a binomial response variable indicating whether a set of responses were tuned (1) or not (0), *Area* : *State* denotes an interaction between brain area and behavioural state, (1 | *Subject*) is a random intercept that takes into account variation between subjects, (1 | *Direction*) is a random intercept that takes into account variation between different directions of motion and (1 | *RF*) is a random intercept that takes into account variation due to the presence or not of a significant receptive field.

To test whether there were statistically significant differences in the tuning strength of neurons between brain areas and behavioural states we used a linear mixed effects (LME) model using the Eq. 5 with the exception that the *ResponseV ariable* term was a continuously valued Tuning Strength variable.

We performed pairwise *F*-tests on model coefficients to test for statistical significance between different areas and behavioural states.

#### Receptive field significance

We used pre-computed p-values for the significance of receptive fields (see Siegle et al., 2019 for details) as provided by the Allen Institute SDK. Briefly, a 2D histogram of spike counts is computed in response to presentations of a 9×9 grid of Gabor stimuli. A chi-square test statistic was then computed as follows:

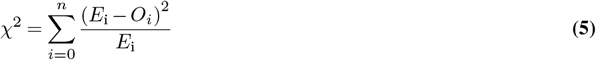

where 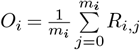 is the observed average response (*R*) of the unit over *m* presentations of the Gabor stimulus at location *i*, and 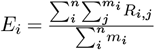 is the expected (grand average) response per stimulus presentation.

The statistical significance of this Chi-square value was then determined by comparing it against null distribution of test statistics calculated after shuffling stimulus locations.

#### Decoding analysis

To decode visual speed from the spike counts of neurons we used a Poisson Independent Decoder (PID) that assumes independent neurons (Zhang et al., 1998; Jazayeri and Movshon, 2006; Graf et al., 2011). For each session, populations of 10 neurons were randomly selected without replacement from individual visual areas. Decoding was conducted separately for each direction of motion.

To compare decoding performance across populations recorded in different sessions, we standardized the number of trials used for decoding by limiting the dataset to 10 trials per stimulus condition (10 trials × 7 speeds = 70 trials total). Leave-one-out cross-validation was used, wherein spike counts from all but one trial per condition (9 trials × 7 speeds = 63 trials) were used to train the decoder, and the remaining trials (1 trial × 7 speeds = 7 trials) were used to test it. This process was repeated 10 times until all 70 trials were tested. Decoder performance was determined as the proportion of correctly classified test trials. For each trial, the predicted visual speed was determined by maximizing the following log-likelihood function:

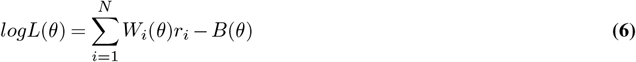

Where *i* indexes over *N* neurons, *W*_*i*_(*θ*) is the log of the mean spike count of neuron *i* for a given stimulus learned from training data, *r*_*i*_ is the number of spikes produced by neuron *i* on the trial being predicted and *B*(*θ*) is a bias correction term calculated as the sum (over N neurons) of mean spike counts for stimulus *θ*.

To test whether there were statistically significant differences between behavioural states for each brain area we first fit an LME with Eq. 5, where the *ResponseV ariable* was Decoding Performance. We then performed pairwise *F*-tests on model coefficients (see analysis of Tuning above).

## Supplementary Figures

**Supplementary Figure 1:**
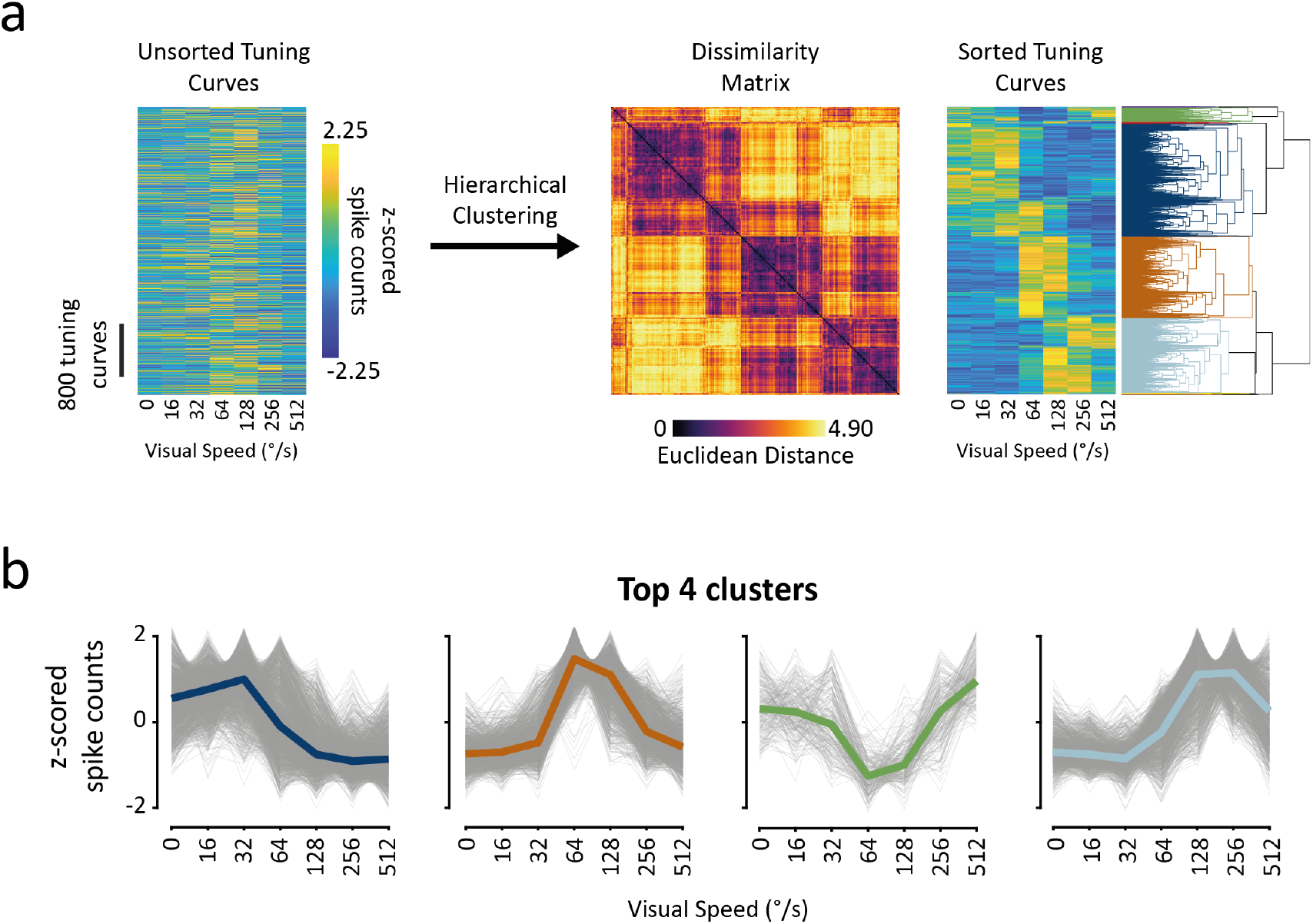
Hierarchical clustering of tuning curves reveals 4 tuning classes. **a** Overview of tuning curve sorting procedure. We used hierarchical sorting to investigate different tuning curve shapes within the mouse visual system. Left: Unsorted tuning curves (z-scored). Middle: dissimilarity matrix after sorting. Right: Sorted tuning curves with aligned dendrogram. **b** Top 4 clusters of tuning curve shape which can be putatively sorted into lowpass, bandpass, bandreject and highpass. Thick coloured traces are medoid of each tuning curve cluster. Thin grey traces are individual tuning curves within a cluster. The same colour code is used as in the dendrogram in a.

**Supplementary Figure 2:**
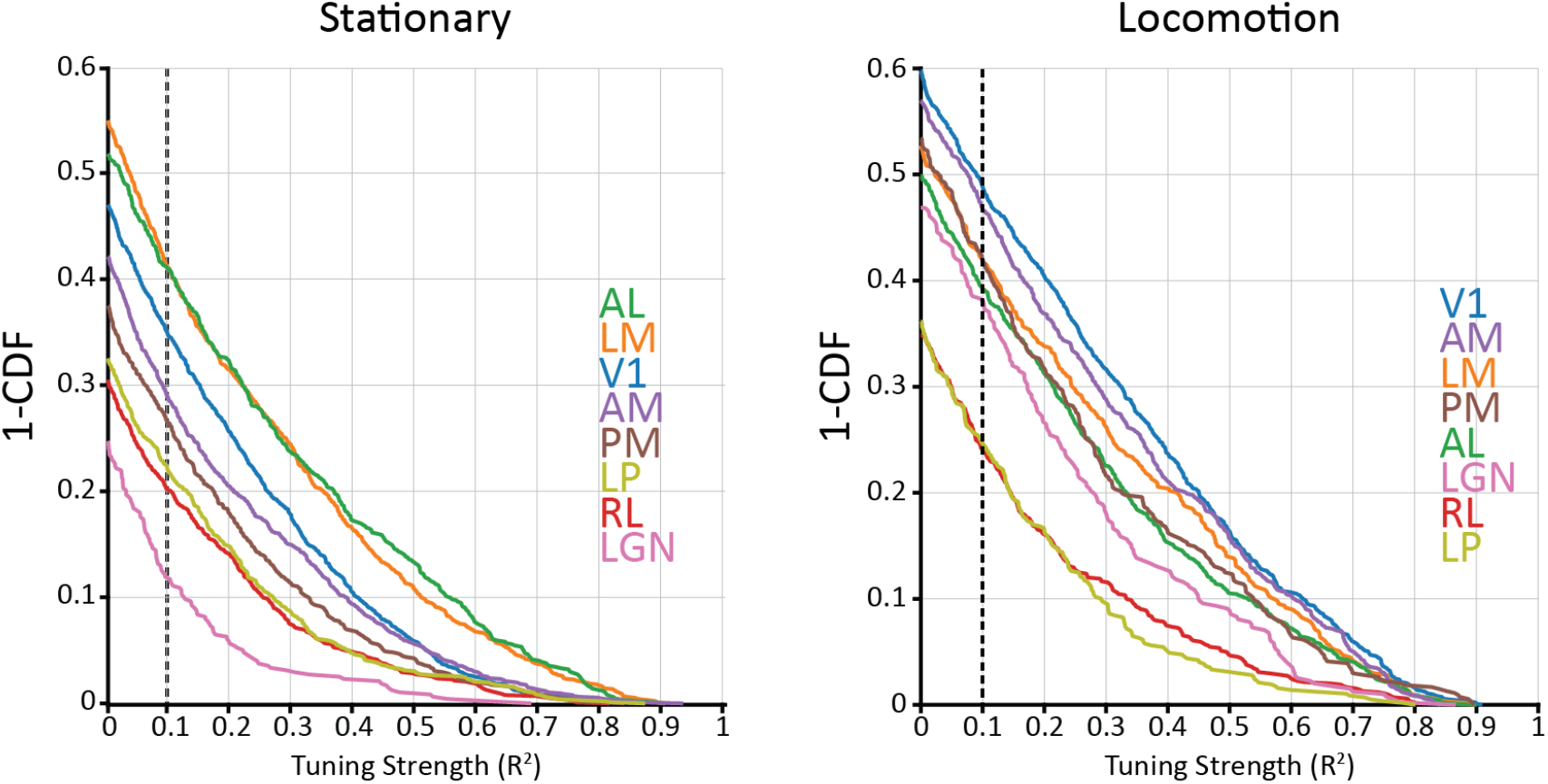
Hierarchical clustering of tuning curves reveals 4 tuning classes. Survival functions (1 - cumulative distribution function) of tuning strength for each visual area during stationary (left) and locomotion (right) states.

